# Hand cold pressor test induces thermogenesis in upper thoracic regions as measured by skin surface infrared thermography

**DOI:** 10.1101/2022.03.22.485388

**Authors:** Natalie Levtova, Emilie-Anne Benoit, Aneet K. Jhajj, Thivaedee Narayana, Dipannita Purkayastha, Ali Salimi, Gabriel Kakon, Meagane E. I. Maurice-Ventouris, Amir Arshiya Kaffash Mohamadi, Tomas Kizhner, Pranamika Khayargoli, Peter J. Darlington

## Abstract

**Background:** Cold exposure may cause health problems and impaired productivity in outdoor or cold-temperature workers. The cold pressor test (CPT) is a laboratory procedure that measures cardiovascular and thermoregulatory responses to acute cold exposure such as metabolic activity in brown adipose tissue. How the body responds to acute cold exposure of a hand is not completely understood. We tested the hypothesis that the upper thorax produces heat during a single hand-CPT, which restores warmth to the cold-expose appendage.

**Objectives:** The objective was to measure skin temperature changes in the upper thoracic regions and the cold-exposed appendage during a CPT. The secondary objective was to determine if cardiovascular or psychological responses during CPT accounted for skin temperature changes.

**Methods:** 50 healthy participants immersed their right hand up to wrist level in 4 °C water for three minutes. Surface skin temperatures were imaged by infrared thermography at baseline, during CPT, and in the recovery phase. Sublingual oral temperature and water bath temperature were recorded throughout the test. Cardiovascular responses were monitored by continuous finger pulse-wave plethysmography. Peak pain and peak stress were reported by the participants on a Likert scale.

**Results:** CPT increased the systolic blood pressure (+22 mmHg, p < 0.001), diastolic blood pressure (+15 mmHg, p < 0.001) and heart rate (+7 beats per minute, p = 0.024). During CPT, skin temperature increased on thoracic regions including mediastinal (+0.5°C, p < 0.021), sternal (+0.5°C, p < 0.002), right supraclavicular (+0.3°C, p < 0.042) and left supraclavicular (+0.3°C, p < 0.016) regions. During CPT, the hand was cooler on ventral (−14.6°C p < 0.001) and dorsal (−15.2°C p < 0.001) sides, and warmed up during recovery. The ventral forearm, dorsal forearm, antecubital fossa, and adjacent medial epicondyle region were significantly cooler throughout the recovery time. The oral temperature did not change during CPT. There were no correlations between the change in mediastinal skin temperature and the sex of the participant, or changes in cardiovascular parameters, peak pain, or peak stress values.

**Conclusions:** Localized hand cooling caused a rapid warming of the thorax, dissipation of cold in the forearm, and rewarming of the hand during recovery. Thermoregulation was not dependant on pain, stress, sex, or cardiovascular changes between participants. By understanding thermoregulation, better approaches can be developed to mitigate the negative impacts of cold exposure.

## Introduction

Extreme temperatures can affect populations with high rates of morbidity such as diabetes and cardiovascular disease (1).Workers exposed to cold were more likely to display increased airway symptoms such as wheeze and cough (2), and long term cold-related circulatory complaints, fatigue, and performance degradation (3). Working in a cold environment was also related to increased reporting of chronic pain (4). Gaining a better understanding of how the body reacts to sudden cold exposure will help manage the implications that extreme climate conditions could have on health. The cold pressor test (CPT) has been used as an indication of blood pressure fluctuations as it relates to meteorological factors such as outdoor temperature (5). Such procedures are done on hands, fingers or feet submerged in a water bath at noxious temperatures (∼0-7°C). CPT has been used to study the autonomic nervous system, thermoregulation and hypertension (11,17,18). When a limb is exposed to cold, a series of reactions occur that serve to preserve the function of the limb and maintain a stable core temperature. The first response occurs prior to cold exposure; the awareness of impending cold produces a small but significant rise in heart rate and blood pressure referred to as the alerting response (6). Localized, acute cold exposure rapidly increases blood viscosity (7) which reduces blood flow (8,9) and perfusion of oxygen into the tissue (10). During CPT, there is a profound increase in sympathetic nerve activity with concomitant parasympathetic withdrawal (11). Temperature sensing receptors in the skin send impulses to the vasomotor center of the brainstem which relays signals to sympathetic nerve pathways. This central neural reflex induces local vasodilation in the cold-affected region and initiates shivering or non-shivering thermogenesis (12). Moreover, it causes vasoconstriction of vascular beds of the kidney (13) and mesenteric (14) circulatory systems. Vasoconstriction of internal blood vessels causes a rise in systolic blood pressure, diastolic blood pressure, systemic vascular resistance, and mean arterial blood pressure (15). Together, these changes help to maintain circulation and temperature control in the cold-affected tissues.

During acute cold exposure, non-shivering thermogenesis occurs in brown adipose tissue. In humans, brown adipose tissue is located subcutaneously at the supraclavicular fossa between anterior neck muscles, axilla, under the clavicles, surrounding the heart, aorta, major arteries, and in pericardial mediastinal fat (16). Acute cold exposure induced brown adipose tissue metabolic activity in thoracic regions, particularly in the right and left supraclavicular fossae (19,20). The overall amounts of brown adipose tissue in the adult human are considered to be rather small, and appear to contribute modestly to the basal metabolic rate (21). However, it was shown using thermal imaging that CPT increased skin temperature of the supraclavicular fossa increases by 0.3–0.7 °C, indicating that there was localized thermogenesis around the brachial arteries which leads to the arms (16,22). This would be essential for thermoregulation because limbs cannot produce significant amounts heat; limbs rely on circulatory control to maintain temperature homeostasis (23,24). One form of circulatory control is counter-current heat exchange, a process whereby venous blood exchanges heat with warm arterial blood thereby attenuating heat loss in the cold-affected limb. The functional anatomy for counter current heat exchange was shown in the legs of wading birds (25,26), however, it has not been studied extensively in humans. Theoretical models predicted that a small but significant amount of heat transfer occurs in the human limb due to counter-current heat exchange in the forearm (24,27).

Infrared thermography is a non-invasive method to assess skin temperature changes. Digital thermal imaging has been recently used to assess febrile temperature (28), bodily maps of emotions (29), and to infer brown adipose activity (16). In the present study, we tested the hypothesis that acute hand cooling induces a rapid temperature increase in the upper thorax. We obtained evidence for circulatory control of arm temperature during CPT using infrared thermography and cardiovascular monitoring.

## Materials and methods

### Participants

This study was approved by the Concordia University human research ethics committee which follows the Declaration of Helsinki principles. Informed signed consent was obtained from 60 healthy participants of age 18 and older. Their health status was determined by self-reporting according to standardized questions throughout the screening phone-interview. To control for menstrual cycle phase, female participants were scheduled to participate during their follicular phase (11). Participants were asked to abstain from caffeine, alcohol, and exercise twelve hours prior to participation, and to not eat at least two hours before the study. Relational ties between participant and experimenter, for example family, partner, or spouse, were avoided given the reported association between empathy and stress-test results (30). Height and weight were obtained in the consultation room, and were used to calculate body mass index using the formula body mass (kg) divided by body height (m) squared. Brachial blood pressure and heart rate were measured in a consultation room after the participant had been sitting for at least 15 minutes (Accutorr Plus V, Medaval, Dublin, Ireland). Participants were excluded from the study if their systolic blood pressure was greater than 140 mmHg or less than 90 mmHg, heart rate was greater than 100 beats per minute or less than 50 beats per minute, or a body mass index greater than 27 kg/m^2^. If these conditions were met, they were directed to see a physician. Of the 60 participants who consented to the study, data from 10 participants were removed because four participants had low blood pressure prior to the test and did not undergo CPT, five participants felt faint or fainted during the CPT and researchers stopped the test, and one participant withdrew their hand from the water mid test. Finally, 50 participants completed the CPT.

### CPT procedure

After consultation, participants moved to the testing room (Table 1). The CPT apparatus was not present when the participant entered the testing room to prevent an alerting response. The chair had a reclining function used in the event of fainting. Their left arm was rested on a chest-height table while the cardiovascular monitor was affixed (Nexfin®; BMEYE, Amsterdam, The Netherlands). The Nexfin is a finger plethysmograph which monitors beat-to-beat arterial blood volume fluctuations with waveform analysis using a finger pressure cuff attached to the contralateral (left) middle finger and calibrated with a heart-level sensor (31). Once the cardiovascular signal stabilized, the baseline was recorded for 15 minutes. In a separate room, the cold bath was prepared. The cold apparatus was a modified picnic cooler with freezer packs and a thermometer affixed to the interior walls of the cooler with industrial grade Velcro. Cold water and crushed ice was added until the bath reached a stable temperature. After the baseline recordings were complete, the cold bath was placed on a platform next to the right side of the chair. At least two minutes elapsed prior to hand immersion. This was considered the alerting response time and it was removed from analysis. At minute 17 the hand immersion commenced. They submerged their right hand to wrist level for 3 minutes in a passive open position; they were instructed to not make a fist. Throughout the baseline CPT and recovery, oral temperature was measured sublingually with a digital oral thermometer. Participants self-reported pain and stress throughout the baseline CPT on a visual analogue scale. A Likert scale was used to determine pain and stress where 0 equalled no pain or stress at all, and 10 equalled the worst possible imaginable pain/stress.

**Table 1.**
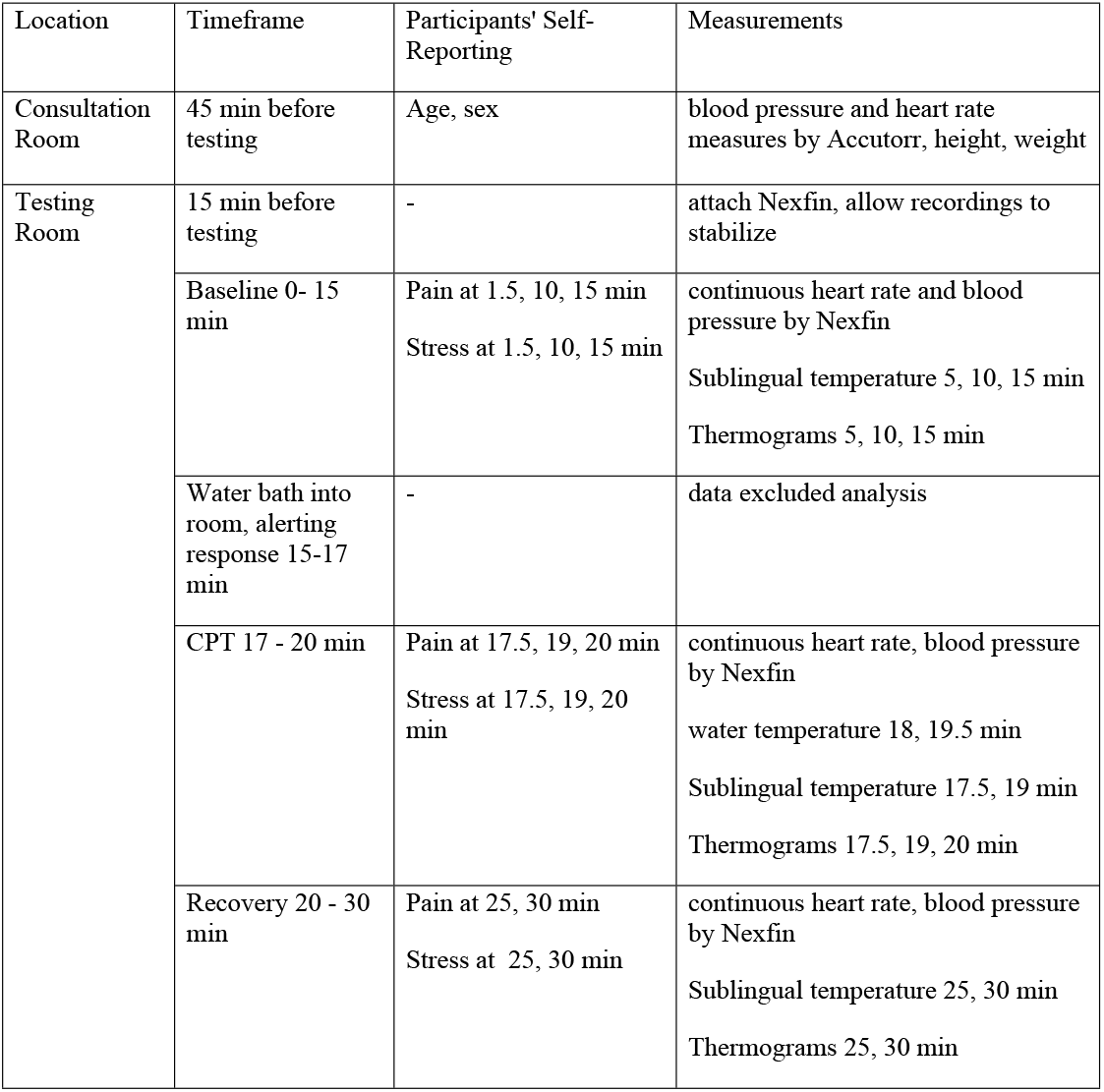
Summary of Study Procedure.

### Thermograms

The FLIR C2 portable infrared camera was used to take thermograms and simultaneous visible-light images (FLIR Systems Inc. Wilsonville, OR, USA). Prior to the test, anatomical landmarks were located and marked on the participant with an erasable pen. To find the jugular notch of the sternum at the top of the midline body structure of the thorax, we followed the clavicle towards the center and a point was placed on the clavicular notch felt at the dip. For the left and right supraclavicular fossae, we placed a point in the angle between the clavicle and the sternocleidomastoid tendon. The thorax image had four regions of interest: middle thorax (mediastinal), upper thorax (sternal, below the jugular notch/manubrium), left supraclavicular fossa and right supraclavicular fossa (between the sternal head of the sternocleidomastoid and the clavicle). For the dorsal hand we marked the intersection between the third extensor digitorum tendon and the purlicue (the fold between the thumb and index finger). For the dorsal forearm we marked halfway between the wrist line and antecubital fossa. For the ventral hand (palmar) we marked the intersection between the purlicue and the center of palm. For the ventral forearm we marked halfway between the wrist line and antecubital fossa. For medial epicondyle, with the participant’s hand in supinated anatomical position (palm up), we marked the medial bony protuberance at the elbow. The regions for the arm in the ventral position (in standard anatomical position) included the antecubital fossa (synonymous with cubital fossa), the adjacent medial epicondyle, forearm (midway between wrist and elbow), and the center of the palm. The dorsal arm regions included the forearm midway between wrist and elbow, and the center dorsum of the hand. Thermograms were taken at a 1 meter distance from the participant at 5 min baseline, 10 min baseline, 15 min baseline, 0.5 min immersion, 2 min immersion, immediately after hand removed from water, 5 min recovery, 10 min recovery (Table 1). The testing room did not contain other infrared light sources. The left arm was not imaged.

### Data analysis

Thermogram files were renamed with randomized codes to ensure that the analysis was done blinded. Two researchers who were unaware of the file codes, independently analyzed the thermograms using FLIR Tools software. Rectangles of 16 × 12 mm were placed on the thoracic regions of interest. For the thermograms of the cold exposed arm, circles diameter 8 mm were placed on the regions of interest. The mean temperature for each region was recorded. The placement of the regions of interest were made using anatomical landmarks visible on the digital picture overlay. For each participant, skin temperature and cardiovascular readings were averaged into three phases: baseline (t1, t2, t3), during CPT (t3.5, t4, t5), and recovery (t6, t7).

Thermograms were excluded from analysis if clothing obstructed the clavicle or mediastinal region, the elbow region or thorax was partially out of frame, or the perpendicular axis of the elbow was on an oblique angle. The number of thermograms analyzed were 50 for right supraclavicular, 49 for left supraclavicular, 50 for sternal, 50 for mediastinal, 49 for ventral hand, 48 for dorsal hand, 45 for ventral forearm, 45 for antecubital fossa, 45 for medial epicondyle region, 43 for and dorsal forearm. The null hypothesis was tested with an analysis of variance (ANOVA, p<0.05) and Tukey’s test or Bonferroni tests. Software used was SPSS Statistics 24.0 (IBM, New York, USA). Pearson’s test was used for correlation analysis.

## Results

### Cardiovascular Response to CPT

Fifty healthy participants completed a three minute CPT by immersing their right hand up to wrist level into ice-cold water. Of the participants who completed the CPT, 29 were female, 21 male, with an average age of 23.9 ± 5.4 years (range 20-43), weight of 68.7 ± 14 kg, and height of 1.70 ± 0.1 m. The participants self-reported their ethnicity as 31 White, six Middle Eastern, five Black, three Asian, two East Asian, one Latin, and two of mixed ethnicity. The average water temperature measured was 2.8 +/- 0.7 °C (N=50). The participants’ systolic blood pressure, diastolic blood pressure, and heart rate increased during CPT and returned towards baseline in the recovery time, while the oral sublingual temperature remained constant throughout the test (Table 2).

**Table 2.** Cardiovascular, oral, and skin temperature measurements in 50 healthy participants undergoing CPT.

| Variable | BL | BL vs. CPT | CPT | CPT vs. REC | REC | BL vs. REC |
| --- | --- | --- | --- | --- | --- | --- |
| systolic BP mmHg | $130 \pm 18$ | *** | $152 \pm 21$ | 0.008 | $139 \pm 19$ | |
| diastolic BP mmHg | $82 \pm 12$ | *** | $97 \pm 15$ | 0.002 | $87 \pm 13$ | |
| heart rate bpm | $70 \pm 10$ | 0.024 | $77 \pm 14$ | 0.003 | $68 \pm 11$ | |
| Oral temp. °C | $36.6 \pm 0.2$ | | $36.5 \pm 0.2$ | | $36.6 \pm 0.2$ | |
| R Supraclavicular °C | $35.4 \pm 0.7$ | 0.042 | $35.7 \pm 0.6$ | | $35.6 \pm 0.8$ | |
| L Supraclavicular °C | $35.1 \pm 0.7$ | 0.016 | $35.5 \pm 0.6$ | | $35.4 \pm 0.6$ | |
| Mediastinal °C | $34.0 \pm 0.8$ | 0.021 | $34.5 \pm 0.8$ | | $34.3 \pm 0.9$ | |
| Sternal °C | 33.9 ± 0.6 | 0.002 | 34.4 ± 0.7 |  | 34.3 ± 0.7 | 0.017 |
| Ventral Hand (Palm) °C | 31.8 ± 2.4 | *** | 17.2 ± 2.7 | *** | 26.2 ± 2.5 | *** |
| Dorsal Hand °C | 30.4 ± 2.1 | *** | 15.2 ± 2.2 | *** | 21.9 ± 1.8 | *** |
| Ventral Forearm °C | 32.7 ± 0.9 | *** | 31.8 ± 1.4 | 0.008 | 31.1 ± 1.0 | *** |
| Dorsal Forearm °C | 31.6 ± 0.9 |  | 31.5 ± 0.1 | *** | 30.5 ± 1.2 | *** |
| Antecubital Fossa °C | 33.7 ± 0.8 |  | 33.9 ± 0.8 |  | 33.0 ± 1.0 | *** |
| Medial Epicondyle °C | 32.3 ± 0.9 |  | 31.9 ± 1.1 |  | 31.4 ± 1.6 | 0.002 |

Cardiovascular and temperature recording during baseline (BL), CPT and recovery (REC). SD is shown. BP = blood pressure. Univariate ANOVA was performed with post hoc Tukey's test for multiple comparisons. Where the differences are significant, P values are shown in columns for baseline as compared to CPT, CPT compared to recovery, and baseline as compared to recovery. The exact p values are indicated, or \*\*\*p<0.001.

### Skin temperature changes

Skin temperatures were measured on the anterior thorax, cold exposed hand, and arm before, during, and after the CPT. We observed skin temperature increases on the thoracic regions during CPT and in the recovery phase (Table 2, Fig 1 a-f). The right and left supraclavicular regions were significantly increased during CPT as compared to baseline. The temperature changes observed on the right and left supraclavicular regions were correlated with each other (Pearson’s correlation of 0.827). The mediastinal and sternal regions were significantly increased during CPT, and the sternal region remained elevated during recovery. With respect to the cold exposed hand, we observed ventral and dorsal skin temperature decreases during CPT which partially rewarmed during recovery (Table 2, Fig 2a-f,). The ventral forearm was cooler immediately after CPT and continued to cool during recovery. The dorsal forearm was cooler during recovery time as compared to CPT or baseline. The antecubital fossa skin temperature was cooler during recovery compared to baseline, and the adjacent medial epicondyle region was cooler during recovery, albeit to a lesser extent than observed for the antecubital fossa (Table 2).

**Fig 1.**
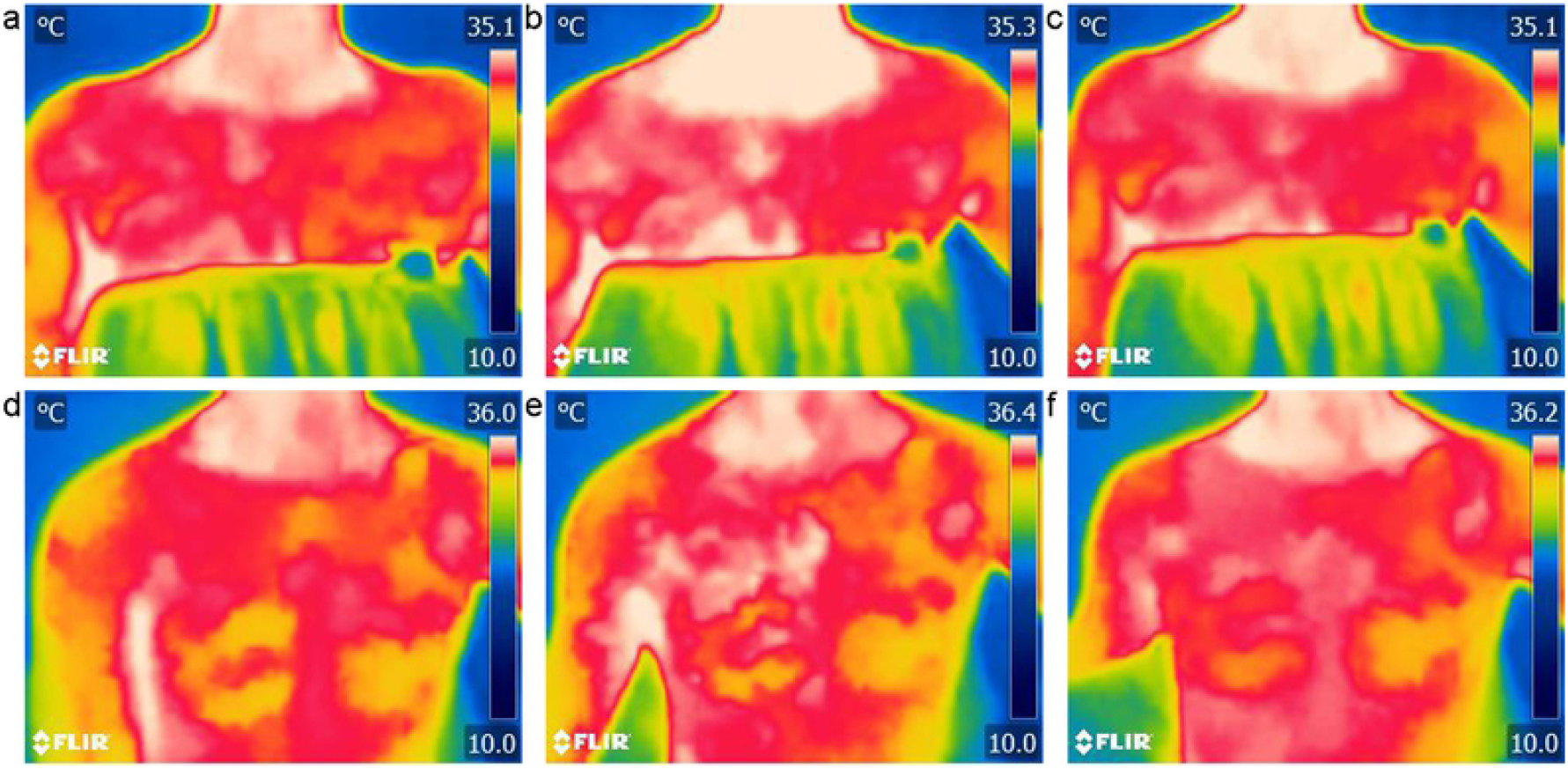
Surface skin temperatures on the anterior thorax. Representative surface skin temperatures of a female participant (top row) and male participant (bottom row) are shown for baseline (a, d), CPT (b, e) and recovery (c, f).

**Fig 2.**
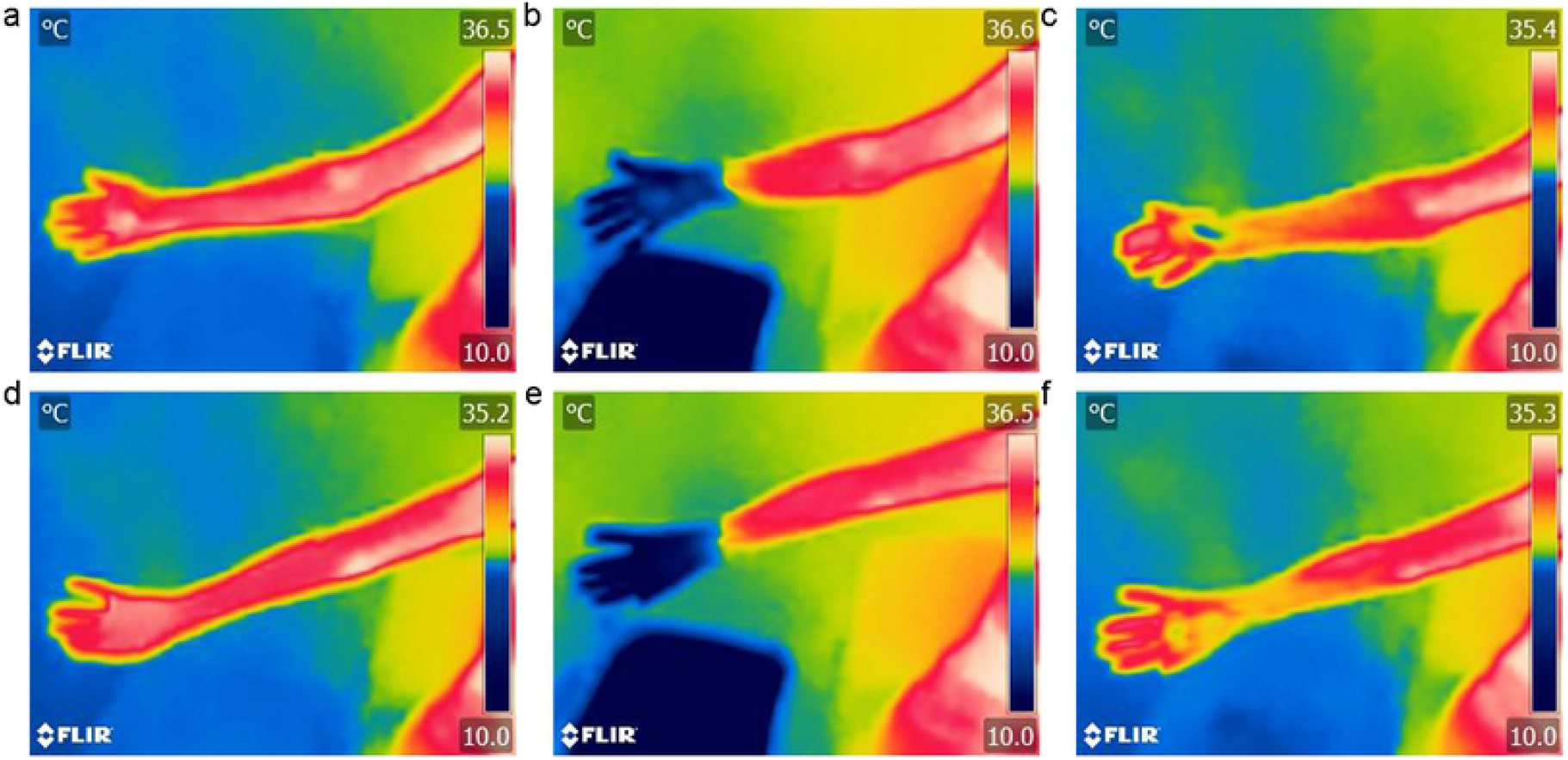
Surface skin temperatures on hand and arm. Surface skin temperatures were measured on the hand and arm in ventral (top row) and dorsal (bottom row) positions. The pictures represent baseline (a, d), CPT (b, e) and recovery (c, f). The thermograms for the CPT was taken immediately after withdrawal of the hand from the ice bath at 3 minutes.

To account for possible psychophysiological variables, the participants were asked to report their pain and stress. The peak pain and peak stress values correlated to each other, however, they did not correlate significantly to the change in mediastinal skin temperature (Table 3). Cardiovascular changes in heart rate and systolic blood pressure did not correlate significantly to the change in mediastinal skin temperature (Table 4). When sex was analyzed as a confounding variable, there were no significant difference in the change such as mediastinal skin temperature during CPT compared between female and male participants (p = 0.94 unpaired t test). Thus, the rapid skin temperature changes did not appear to be influenced by self-reported pain, stress, cardiovascular measures, or sex. In summary, a three minute hand CPT induced rapid warming of the thorax, a sharp drop in temperature of the hand with subsequent rewarming, and a dissipation of cold throughout the forearm during recovery.

**Table 3.** Pearson correlations between change in psychological variables with change in mediastinal skin temperature during CPT.

| Pearson correlations | peak pain | peak stress | $\Delta$ mediastinal skin temperature |
| --- | --- | --- | --- |
| peak pain | 1 | 0.628 ** | -0.120 |
| peak stress | 0.628 ** | 1 | 0.084 |
| $\Delta$ mediastinal skin temperature | -0.120 | 0.084 | 1 |

\*\* $p < 0.01$ significance level, $\Delta$ = change between CPT and baseline

**Table 4.** Pearson correlations between change in cardiovascular variables with change in mediastinal skin temperature during CPT.

| Pearson correlations | $\Delta$ heart rate | $\Delta$ systolic blood pressure | $\Delta$ mediastinal skin temperature |
| --- | --- | --- | --- |
| $\Delta$ heart rate | 1 | 0.249 | 0.001 |
| $\Delta$ systolic blood pressure | 0.249 | 1 | 0.209 |
| $\Delta$ Mediastinal Skin Temperature | 0.001 | 0.209 | 1 |

No significant correlations were observed. $\Delta$ = change between CPT and baseline

## Discussion

Acute cold exposure causes a rapid cardiovascular and thermal response that protects the exposed limb from hypothermia and maintains a constant core temperature. Our raw data is found in a supplemental data file (S1 File). We observed significant increases in systolic blood pressure, diastolic blood pressure, and heart rate which is the expected result of CPT (32). Furthermore, we observed the expected temperature increases the thoracic regions including right and left supraclavicular fossa. These results are similar to those reported by Sacks and Symonds 2013, who collected thermograms from a small sample of people undergoing CPT (16). Their analysis focused on skin temperature changes in the supraclavicular regions where brown adipose tissue is known to exist. Brown adipose tissue can induce thermogenesis in these supraclavicular regions due to increased metabolic activity as shown with positron emission scanning of metabolic tracers (33). We also detected skin temperature increases on the central mediastinal and upper sternal areas of the thoracic midline. Mediastinal skin temperature on the thorax may be influenced by dilation of epicardia coronary arteries in response to an increased myocardial metabolic demand during CPT. This would increase blood flow, and therefor increase skin temperature in the upper thoracic region (34). Another explanation is activity of deep muscles in the cervico-thoracic region that can produce heat by non-shivering thermogenesis (20). Thermogenesis is also potentially caused by brown or beige adipose tissue in the thorax. Beige adipose is a unique blend of brown adipose tissue and white adipose tissue that can have metabolic activity (35,36). One or all of these mechanisms could have contributed to the skin temperature increases we observed on the thorax during exposure of the hand to acute cold.

The sublingual temperature did not change during CPT or in recovery, which is consistent with other findings; short-term cold exposure of a limb did not alter body core temperature (37). Core temperature does not change during acute localized hand-cooling, as was shown previously using an intestinal thermometer during 3 minute hand-CPT in a normothermic ambient environment (38). It takes over 10-15 minutes of pressor response for core temperatures to show a difference which is longer than our three minute exposure protocol (39). As expected, the temperature of the cold-exposed hand decreased substantially, and partially rewarmed during the 10 minute recovery. In the cold-exposed forearm, which was not submerged in the ice bath, we observed modest skin temperature decreases immediately after CPT and during the recovery time. That finding indicates that venous return is carrying cool blood away from the cold-exposed hand. As warm arterial blood has the potential to exchange heat with the cooler venous return, the forearm may represent a form of counter-current heat exchange that maintains core temperature and slowly rewarms the hand during acute cold exposure.

We investigated the possible link between thermogenesis and the psychological factors of pain and stress. Psychological factors can influence cardiovascular changes during CPT, for example, we recently reported a positive correlation with the amount of pain reported by the participants and their change in heart rate (40). Other researchers showed that the emotion of anxiety raised thoracic skin temperature using psychological testing and body temperature mapping (29). In the present study, we did not observe correlations between self-reported peak pain or peak stress with the change in mediastinal skin temperature. Furthermore, we did not find correlations between cardiovascular changes and the change in mediastinal skin temperature. Together, these data suggest that thermogenesis during CPT is a rapid neural or hormonal reflex that is unaffected by pain, stress, sex, or cardiovascular factors.

The overarching purpose of this study was to learn more about the effects of acute cold exposure on human physiology. The technical limitations of our study included the use of an industrial grade thermal camera which has less resolution and precision than clinical grade thermal cameras. Some thermograms had to be excluded due to slightly incorrect positioning of the arm. Having a fixed camera and giving the participants better instructions on arm positioning would correct these limitations. One of the advantages of using a portable thermal camera is its low cost and flexibility. A portable thermal camera could be used for field work aimed at capturing the thermal responses in people working in a cold environment. Our study validates the use of thermal imaging as a non-invasive, cost-effective and flexible method to study thermogenesis. By understanding the fundamental mechanisms of action that the body uses to maintain thermoregulation, we can better guide new approaches to prevent cold exposure health issues.

## References

1. Gronlund CJ, Sullivan KP, Kefelegn Y, Cameron L, O’Neill MS. Climate change and temperature extremes: A review of heat- and cold-related morbidity and mortality concerns of municipalities. Maturitas. 2018 Aug;114:54–9.

2. Stjernbrandt A, Stenfors N, Liljelind I. Occupational cold exposure is associated with increased reporting of airway symptoms. Int Arch Occup Environ Health. 2021 Nov;94(8):1945– 52.

3. Auttanate N, Chotiphan C, Maruo SJ, Näyhä S, Jussila K, Rissanen S, et al. Cold-related symptoms and performance degradation among Thai poultry industry workers with reference to vulnerable groups: a cross-sectional study. BMC Public Health. 2020 Dec;20(1):1357.

4. Farbu EH, Skandfer M, Nielsen C, Brenn T, Stubhaug A, Höper AC. Working in a cold environment, feeling cold at work and chronic pain: a cross-sectional analysis of the Tromsø Study. BMJ Open. 2019 Nov;9(11):e031248.

5. Ni Y, Miao Q, Zheng R, Miao Y, Zhang X, Zhu Y. Individual sensitivity of cold pressor, environmental meteorological factors associated with blood pressure and its fluctuation. Int J Biometeorol. 2020 Sep;64(9):1509–17.

6. Lacey JI, Lacey BC. The law of initial value in the longitudinal study of autonomic constitution: reproducibility of autonomic responses and response patterns over a four-year interval. Ann N Y Acad Sci. 1962;98(4):1257–90.

7. Çinar Y, Şenyol AM, Duman K. Blood viscosity and blood pressure: role of temperature and hyperglycemia. American journal of hypertension. 2001;14(5):433–8.

8. Houben H, Thien T, Wijnands G, Van’t Laar A. Effects of cold exposure on blood pressure, heart rate and forearm blood flow in normotensives during selective and non-selective beta-adrenoceptor blockade. BrJClinPharmacol. 1982;14(6):867–70.

9. Wong F, Logan A, Blendis L. The effect of varying sodium intake on blood volume, forearm blood flow and vascular responsiveness to sympathetic stimulation in pre-ascitic cirrhosis. ClinInvestMed. 1996 Jun;19(3):184–94.

10. Novikova IN, Dunaev AV, Sidorov VV, Krupatkin AI. [Functional Status of Microcirculatory-Tissue Systems during the Cold Pressor Test]. Fiziol Cheloveka. 2015 Dec;41(6):95–103.

11. Wirch JL, Wolfe LA, Weissgerber TL, Davies GAL. Cold pressor test protocol to evaluate cardiac autonomic function. Appl Physiol Nutr Metab. 2006 Jun;31(3):235–43.

12. Morrison SF. Central neural control of thermoregulation and brown adipose tissue. Auton Neurosci. 2016 Apr;196:14–24.

13. Boddi M, Sacchi S, Lammel RM, Mohseni R, Serneri GG. Age-related and vasomotor stimuli-induced changes in renal vascular resistance detected by Doppler ultrasound. Am J Hypertens. 1996 May;9(5):461–6.

14. Chaudhuri KR, Thomaides T, Hernandez P, Alam M, Mathias CJ. Noninvasive quantification of superior mesenteric artery blood flow during sympathoneural activation in normal subjects. Clin Auton Res. 1991 Mar;1(1):37–42.

15. Fagius J, Karhuvaara S, Sundlof G. The cold pressor test: effects on sympathetic nerve activity in human muscle and skin nerve fascicles. Acta Physiologica. 1989;137(3):325–34.

16. Sacks H, Symonds ME. Anatomical locations of human brown adipose tissue: functional relevance and implications in obesity and type 2 diabetes. Diabetes. 2013 Jun;62(6):1783–90.

17. Hines EA, Brown GE. The cold pressor test for measuring the reactibility of the blood pressure: data concerning 571 normal and hypertensive subjects. AmHeart J. 1936;11(1):1–9.

18. Victor RG, Leimbach WN, Seals DR, Wallin BG, Mark AL. Effects of the cold pressor test on muscle sympathetic nerve activity in humans. Hypertension. 1987 May;9(5):429–36.

19. Orava J, Nuutila P, Lidell ME, Oikonen V, Noponen T, Viljanen T, et al. Different metabolic responses of human brown adipose tissue to activation by cold and insulin. Cell Metab. 2011 Aug 3;14(2):272–9.

20. U Din M, Raiko J, Saari T, Kudomi N, Tolvanen T, Oikonen V, et al. Human brown adipose tissue [(15)O]O2 PET imaging in the presence and absence of cold stimulus. Eur J Nucl Med Mol Imaging. 2016 Sep;43(10):1878–86.

21. Chechi K, Carpentier AC, Richard D. Understanding the brown adipocyte as a contributor to energy homeostasis. Trends Endocrinol Metab. 2013 Aug;24(8):408–20.

22. Symonds ME, Henderson K, Elvidge L, Bosman C, Sharkey D, Perkins AC, et al. Thermal imaging to assess age-related changes of skin temperature within the supraclavicular region co-locating with brown adipose tissue in healthy children. J Pediatr. 2012 Nov;161(5):892–8.

23. Huttunen P, Hirvonen J, Kinnula V. The occurrence of brown adipose tissue in outdoor workers. Eur J Appl Physiol Occup Physiol. 1981;46(4):339–45.

24. Mitchell JW, Myers GE. An analytical model of the counter-current heat exchange phenomena. Biophys J. 1968 Aug;8(8):897–911.

25. Midtgård U, Bech C. Responses to catecholamines and nerve stimulation of the perfused rete tibiotarsale and associated blood vessels in the hind limb of the Mallard (Anas platyrhynchos). Acta Physiol Scand. 1981 May;112(1):77–81.

26. Fitzpatrick MJ, Mathewson PD, Porter WP. Validation of a Mechanistic Model for Non-Invasive Study of Ecological Energetics in an Endangered Wading Bird with Counter-Current Heat Exchange in its Legs. PLoS One. 2015;10(8):e0136677.

27. Barnett SS, Smolinski P, Vorp DA. A Three-Dimensional Finite Element Analysis of Heat Transfer in the Forearm. Comput Methods Biomech Biomed Engin. 2000;3(4):287–96.

28. Ogawa T, Tsukuda Y, Suzuki Y, Hiratsuka S, Inoue R, Iwasaki N. Utility of thermal image scanning in screening for febrile patients in cold climates. Journal of Orthopaedic Science. 2021 Sep;S0949265821002682.

29. Nummenmaa L, Glerean E, Hari R, Hietanen JK. Bodily maps of emotions. Proceedings of the National Academy of Sciences. 2014 Jan 14;111(2):646–51.

30. Phillips AC, Carroll D, Hunt K, Der G. The effects of the spontaneous presence of a spouse/partner and others on cardiovascular reactions to an acute psychological challenge. Psychophysiology. 2006 Nov;43(6):633–40.

31. Sipkens LM, Treskes K, Ariese-Beldman K, Veerman DP, Boer C. Application of Nexfin noninvasive beat-to-beat arterial blood pressure monitoring in autonomic function testing. Blood Press Monit. 2011 Oct;16(5):246–51.

32. Youssef M, Ghassemi A, Carvajal Gonczi CM, Kugathasan TA, Kilgour RD, Darlington PJ. Low Baseline Sympathetic Tone Correlates to a Greater Blood Pressure Change in the Cold Pressor Test. AerospMedHumPerform. 2018 Jun 1;89(6):503–9.

33. Virtanen KA, Lidell ME, Orava J, Heglind M, Westergren R, Niemi T, et al. Functional brown adipose tissue in healthy adults. N Engl J Med. 2009 Apr 9;360(15):1518–25.

34. Nitenberg A, Chemla D, Antony I. Epicardial coronary artery constriction to cold pressor test is predictive of cardiovascular events in hypertensive patients with angiographically normal coronary arteries and without other major coronary risk factor. Atherosclerosis. 2004;173(1):115–23.

35. Wu J, Boström P, Sparks LM, Ye L, Choi JH, Giang A-H, et al. Beige adipocytes are a distinct type of thermogenic fat cell in mouse and human. Cell. 2012 Jul 20;150(2):366–76.

36. Jespersen NZ, Larsen TJ, Peijs L, Daugaard S, Homøe P, Loft A, et al. A classical brown adipose tissue mRNA signature partly overlaps with brite in the supraclavicular region of adult humans. Cell Metab. 2013 May 7;17(5):798–805.

37. Daanen H a. M. Finger cold-induced vasodilation: a review. Eur J Appl Physiol. 2003 Jun;89(5):411–26.

38. Cui J, Shibasaki M, Low DA, Keller DM, Davis SL, Crandall CG. Heat stress attenuates the increase in arterial blood pressure during the cold pressor test. JApplPhysiol (1985). 2010 Nov;109(5):1354–9.

39. Sendowski I, Savourey G, Launay J-C, Besnard Y, Cottet-Emard J-M, Pequignot J-M, et al. Sympathetic stimulation induced by hand cooling alters cold-induced vasodilatation in humans. EurJApplPhysiol. 2000;81(4):303–9.

40. Kakon G, Mohamadi A-AK, Levtova N, Maurice-Ventouris MEI, Benoit E-A, Chouchou F, et al. Elevated Heart Rate and Pain During a Cold Pressor Test Correlates to Pain Catastrophizing. Appl Psychophysiol Biofeedback. 2021 Dec;46(4):359–66.

